# Evaluating the Role of *Anopheles* Mosquitoes in the Global Spread of Arboviruses: A Review of Laboratory-Confirmed Viral Competence

**DOI:** 10.1101/2025.10.22.683863

**Authors:** Rosheen S. Mthawanji, Matthew Baylis, Maya Wardeh, Marcus SC Blagrove

## Abstract

Mosquito-borne diseases are a major global health concern, infecting up to 700 million people annually and causing over one million deaths. Of the several genera of biting mosquito, species of *Anopheles* are mostly studied for their ability (vector competence) to transmit *Plasmodium* protozoan parasites, some species of which cause malaria. More than 480 species of *Anopheles* have been described worldwide and about 70 of these are responsible for *Plasmodium spp*. transmission. However, the focus on *Anopheles* as vectors of *Plasmodium* has led to a relative lack of study about the ability of *Anopheles* to transmit viruses.

Some *Anopheles* species have been experimentally confirmed as competentfor various arboviruses. In most cases they are secondary vectors, with relatively low competence, contributing to overall transmission while other species of mosquito or other vector are responsible for sustained transmission. Although secondary vectors may contribute less to transmission, they may play important epidemiological roles by extending transmission seasons and/or providing a means of virus overwintering.

Here, we conducted an extensive review of scientific repositories, accessing a diverse range of journals and reports to build an exhaustive database of known *Anopheles* competence for arboviruses. Opinion pieces, modelling papers lacking experimental confirmation, and papers without vector competence data were excluded. In total, we reviewed 7,343 papers, and after exclusions, we retained 427 papers.

We analysed these data, cross-referenced with ability to undergo diapause and geographical range. Our results show that some *Anopheles* spp. have the potential to be secondary vectors of some arboviruses, as well as undergoing diapause as adults in temperate regions of their range. Hence, we infer they could also play an epidemiologically significant role in the overwintering of mosquito-borne viruses.

**Author Summary:** Mosquito-borne diseases remain a major global health challenge, causing over one million deaths annually. Though *Anopheles* mosquitoes are widely studied for their spread of parasites that cause malaria, much less is known about their interaction with viruses. In this study we reviewed scientific articles to identify their role in the spread of mosquito-borne viruses. We examined studies that showed experimental evidence of transmission in the laboratory, and thus could potentially contribute to transmission as secondary vectors. Additionally, some *Anopheles* species in temperate regions can survive winter in a dormant state (diapause) as adults, which could enable viruses to persist between transmission seasons. Together, these results suggest that *Anopheles* mosquitoes may play a larger role in virus transmission than is currently appreciated, especially in temperate regions.

## Introduction

Mosquito-borne viruses present a global health threat due to their broad host range, infecting humans and animals annually. This poses a challenge for disease control experts as the inevitable spread to new locations and hosts complicate understanding vector dynamics (1). During an arbovirus outbreak, the primary vector tends to be the focus of control programmes, whilst secondary vectors may remain under-characterized and insufficiently investigated (2). An understanding of the importance of secondary vectors prior to outbreaks would increase control method efficiency, redirect target response and mitigation procedures and anticipate transmission in regions or seasons where primary vectors are absent or inactive.

Among the hundreds of known arboviruses, only approximately 30 are known to cause human diseases. The Flaviviridae and Togaviridae viral families, particularly Japanese encephalitis (JEV), West Nile virus (WNV), dengue virus (DENV), and Zika virus (ZIKV), are commonly studied for causing severe clinical diseases. Alphaviruses in the Togaviridae family, such as Chikungunya (CHIKV), the encephalitic viruses (Eastern, Western, and Venezuelan), and the Ross River virus (RRV), are also of great public health importance(3). Arbovirus transmission requires a competent vector: one capable of becoming infected and having the capability to then transmit the virus from one host to another. Important mosquito vectors include *Aedes aegypti, Aedes albopictus*, and *Culex quinquefasciatus*.; these species are very widely distributed and are competent vectors of several arboviruses (2,5) but many other species are also known to be competent. However, many other species are also capable of transmitting viruses, and understanding the full range of vector competence is essential for effective disease control (4).

In tropical countries arbovirus transmission tends to be year-round. For example, dengue and chikungunya virus transmission is sustained through mosquitoes being active all year, even though mosquito populations decline in the colder months (6). In this situation, competent secondary vectors could add to the overall burden, but the primary vectors are sufficient on their own to sustain transmission. However, different trends are seen in temperate regions, with viruses like West Nile virus (WNV) and Usutu virus (USUV), as mosquitoes are only active for a smaller portion of the year (7). There is an urgent need to understand the dynamics and methods around how the viruses persist in cold temperatures, when mosquitoes are not active. These may include vertical transmission (9) within mosquito populations, persistent infection in vertebrate hosts (6), or survival of infected mosquitoes through cold seasons. The evidence for long-term vertebrate infection for most arboviruses is limited, and vertical transmission has been confirmed for only a few arboviruses. The possibility that infected mosquitoes, particularly secondary vectors, could overwinter and reinitiate transmission in the spring remains underexplored.

Overwintering refers to the survival of organisms through adverse seasonal conditions, often via behavioural or physiological adaptations. In mosquitoes, this may involve diapause, a form of dormancy that allows survival through cold or dry periods. For example, many *Anopheles* mosquitoes diapause during temperature extremes. *Anopheles* mosquitoes use diapause to persist in long dry seasons in sub-Saharan Africa; instead of using their resources for reproduction, most processes halt and are allocated to longevity (11). *Anopheles colluzzi* are known to diapause in harsh dry conditions (11) while *Anopheles freeborni* diapauses in extremely low temperatures (12). Both species diapause in their adult forms.

The UK is a potential example of where virus may persist in diapausing adults, as the same strain of USUV has persisted in the country for multiple years, despite the transmission season being only during the peak of summer.

Here, we demonstrate that *Anopheles* spp could contribute to overwintering by diapausing as infected (and ultimately infectious) adults. Even low levels of competence in a diapausing *Anopheles spp* could significantly add to the risk of overwintering. Notably, African *Anopheles gambiae* and the Indo-Iranian *Anopheles stephensi* are well-studied due to the malaria-associated burdens in their respective regions (19). To date, only one arbovirus, the O’nyong-nyong virus, is known to be primarily transmitted by *Anopheles* mosquitoes in the field (*Anopheles funestus and An. gambiae*), although other *Anopheles* species have demonstrated the capability to be vectors of alphaviruses in laboratory settings and hence may be secondary vectors (19,20). Understanding which mosquitoes are susceptible or capable of transmitting arboviruses is crucial for controlling threats to human and animal health.

To assess this possibility, we conducted a systematic review of existing literature. Systematic reviews provide a transparent and reproducible method for synthesizing evidence across diverse studies and are particularly valuable when addressing under-researched or fragmented topics.

Here we do this for:

1. laboratory *Anopheles* vector competence, and
2. *Anopheles* diapausing habits to identify viruses in which overwintering in *Anopheles* potentially occurs.

Our aim is to identify arboviruses for which overwintering in *Anopheles* may be plausible, thereby highlighting potential secondary vector roles that may influence long-term virus persistence and seasonal transmission dynamics.

## Methods

### Data Collection

We conducted an extensive review of PubMed, accessing a diverse range of books, journals, and reports. Our systematic search employed multiple keywords:

> “experimental transmission” OR “vector competence” OR “vector-competence” OR refractory OR “infection rate” OR “infection rates” OR “transmission rate” OR “transmission rates” OR “dissemination rate” OR “dissemination rates” AND mosquito

The searches were limited to the English language “Mosquito” was included in all searches. The review covered studies published between 1972 and 2023.

### Exclusion/Inclusion criteria

Duplicate records, systematic reviews, opinion pieces, modelling papers lacking experimental confirmation, and papers without vector competence data were excluded. The inclusion/exclusion criteria ensured that there were no confounding factors favouring virus infection and it was limited to laboratory studies and did not include inferred or assumed competence from field-only data

Information extracted from the papers included the PubMed ID, title, digital object identifier (DOI), the virus of interest with its characteristics (virus class, name, tax ID, lineage, strain, accession number, and culture), mosquito details (species, tax ID, origin, location, lab colony or field-collected, generation, and collection coordinates), and experimental parameters (temperature, infectious dosage, unit used, feeding method, days post-infection, total infected, infection method, and transmission method). Experimental results, including the number infected, infection rate (proportion of mosquitoes with detectable virus after exposure), dissemination rate (proportion of infected mosquitoes in which the virus spread beyond the midgut), and transmission rate (proportion of mosquitoes with virus present in saliva or capable of transmitting it), were also recorded.

#### Screening and Data Verification

We conducted a systematic review to assess the role of *Anopheles* mosquitoes in arbovirus transmission. A total of 7,343 articles were identified with our search term. After removing records based on our above exclusion criteria, this screening yielded 427 articles reporting vector competence in *Anopheles* and other mosquito genera. All data were independently cross-verified independently by two reviewers to ensure consistency and accuracy. The studies collectively covered 95 countries and involved 67 different viruses. For the purposes of this study, we extracted and analysed only the data related to *Anopheles* mosquitoes; all of which are included in the supplementary materials. These data are part of a larger unpublished curated dataset compiling vector competence information across mosquito genera. The full process, including article selection and filtering criteria, is summarized in Figure 1

**Figure 1.**
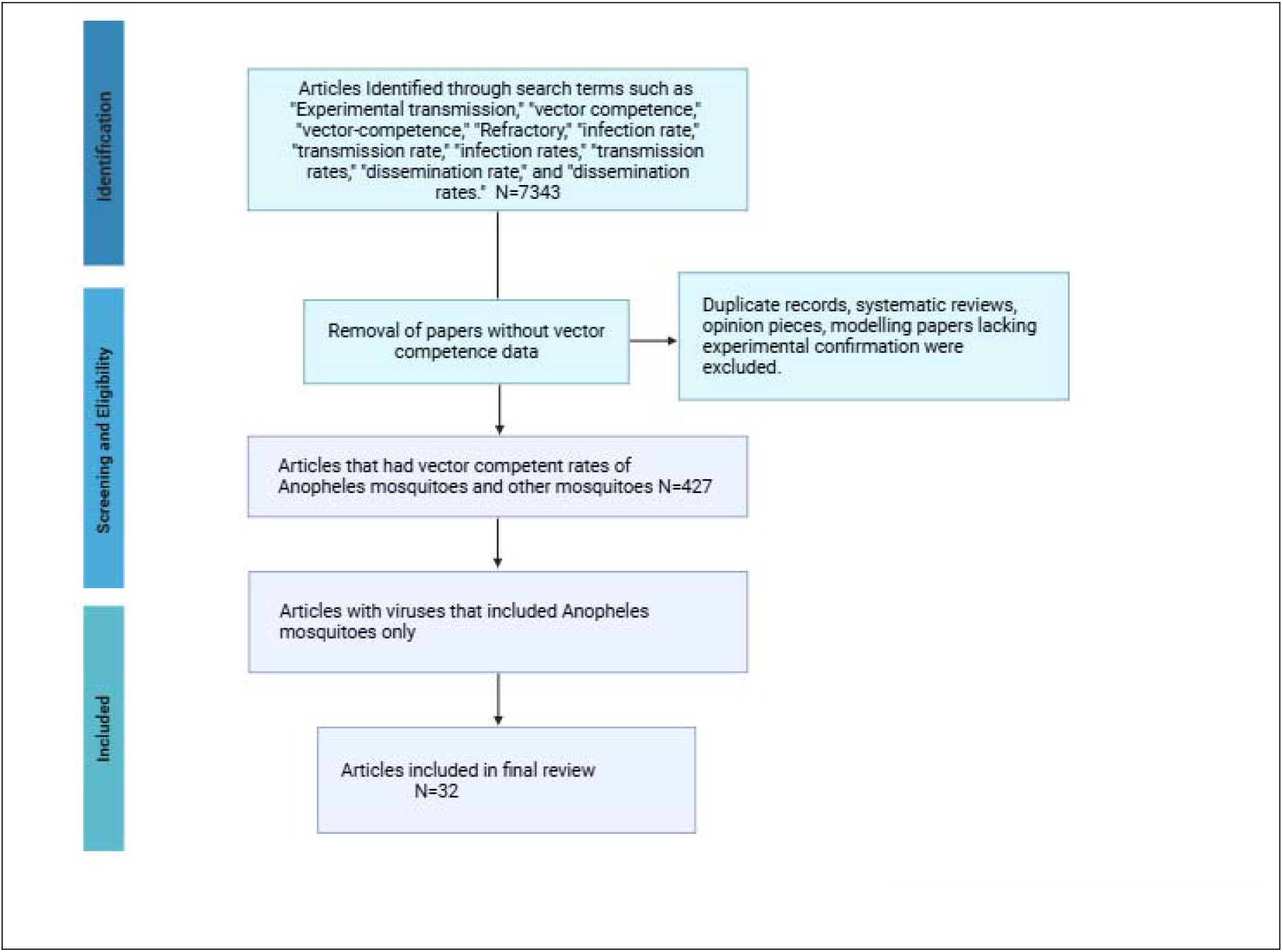
PRISMA Flowchart of main search strategy and article selection for paper selection.

## Results

The vector competence data summarised in Table **1** **shows findings from 32 studies examining** *Anopheles* mosquitoes experimentally exposed to arboviruses in the laboratory. The studies were from 15 countries and presented five virus classes; alphavirus, orthoflavivirus, phlebovirus, orthobunyavirus, and alphamesonivirus and involves **15 *Anopheles* species**, including *An. gambiae, An. stephensi, An. albimanus, An. quadrimaculatus*, and others. For each mosquito-virus pairing, we recorded data on infection, dissemination, and transmission. In cases where part of the experiment was not performed, it is registered as not done in the table (e.g. dissemination but not transmission was tested).

**Table 1.**
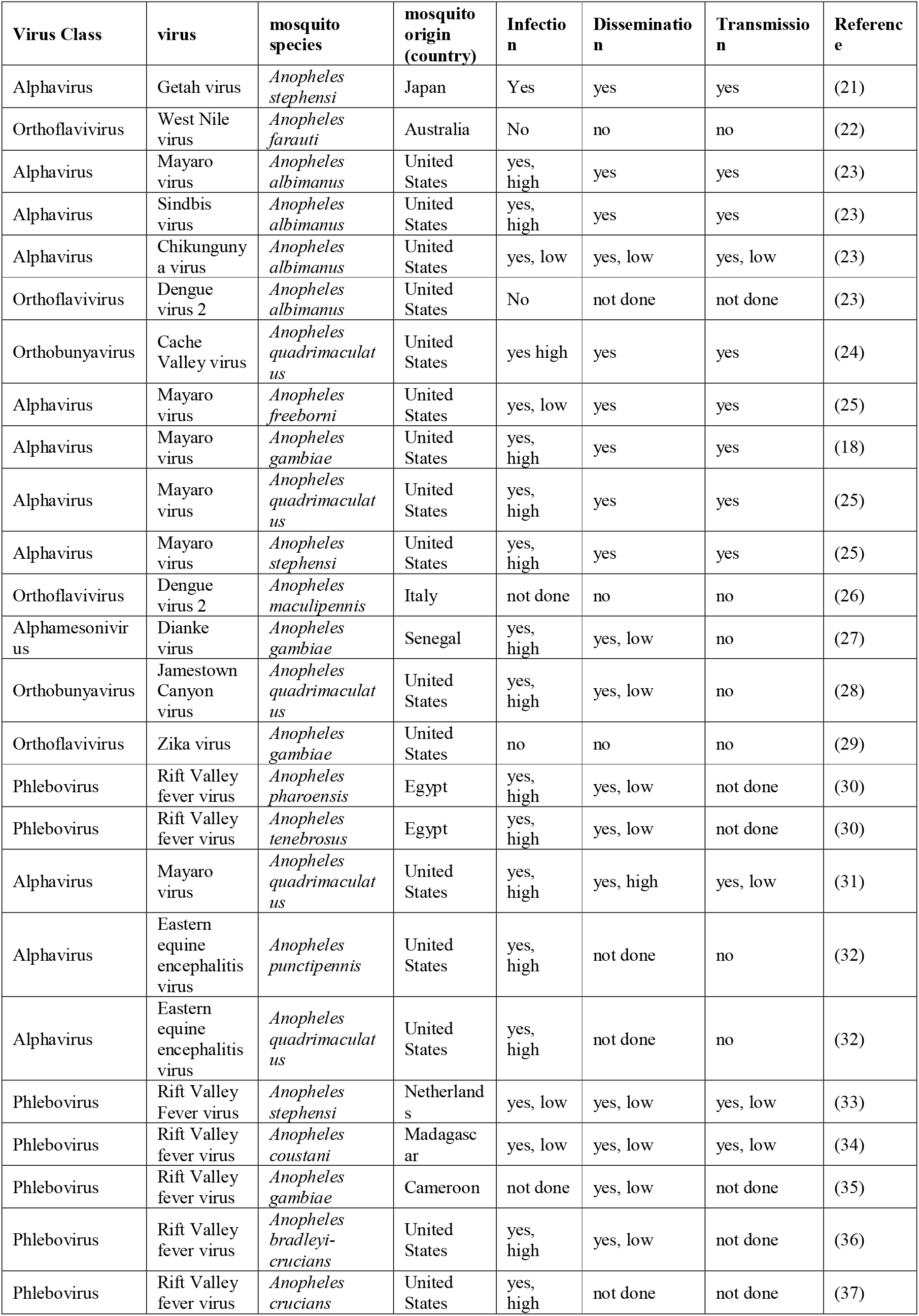
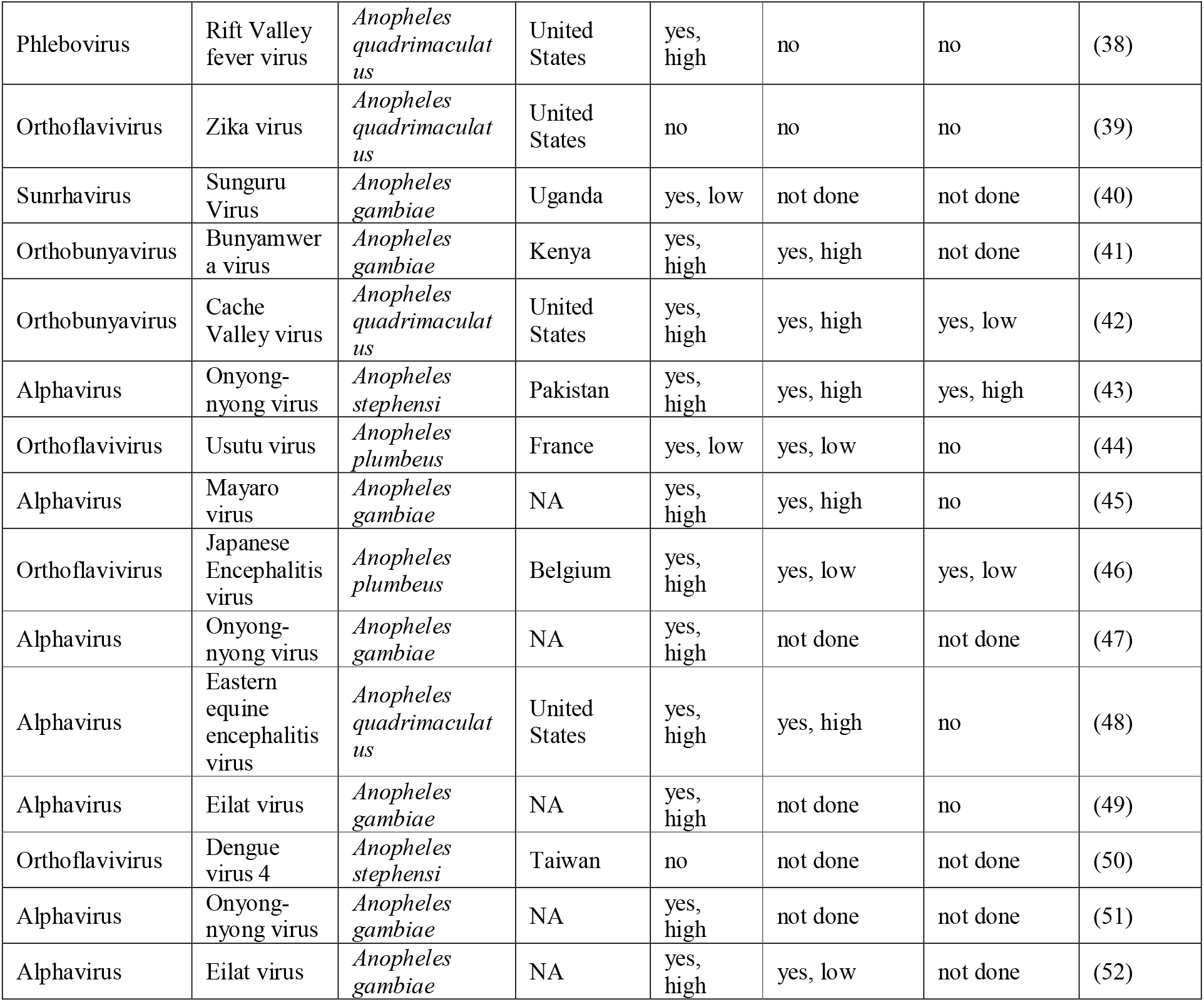
Data extracted from the *Anopheles’* data showing virus classification, mosquito species, Country of origin, vector competence and references. (supplementary material), low percentage values were defined as those ranging from 0% to 40%, whereas high percentage values were defined as those from 50% and above.

The results highlight variation in vector competence across virus families and mosquito species. Notably, several Anopheles species demonstrated high competence for alphaviruses such as Mayaro virus and O’nyong-nyong virus (ONNV), with consistently high infection, dissemination, and transmission rates. In comparison, most Anopheles species showed no competence for orthoflaviviruses such as dengue virus, Zika virus, and West Nile virus, which frequently yielded no infection or transmission. Some species, particularly *An. quadrimaculatus, An. stephensi, and An. gambiae*, also showed susceptibility to selected orthobunyaviruses (e.g., Cache Valley virus, CVV) and phleboviruses (e.g., Rift Valley fever virus, RVFV), although transmission rates varied.

It is important to note that a majority of the studies were conducted in the United States, reflecting a geographic bias in the available data and underscoring the urgent need for a deeper global investigation particularly in endemic or emerging regions where *Anopheles* species are present. Altogether, this dataset shows the potential for *Anopheles* mosquitoes to act as secondary vectors for select arboviruses especially in regions or seasons where primary vectors may be absent, inactive, not year-round, or less abundant

To explore experimental vector-virus interactions in Anopheles mosquitoes, we mapped the viruses and mosquito species included in our dataset onto their respective phylogenies. Figure 2 shows the relationships between arboviruses and Anopheles mosquitoes tested for vector competence in published experimental studies. The results show multiple viruses including Mayaro, Chikungunya, O’nyong-nyong, Sindbis, and Eastern equine encephalitis viruses.

**Figure 2.**
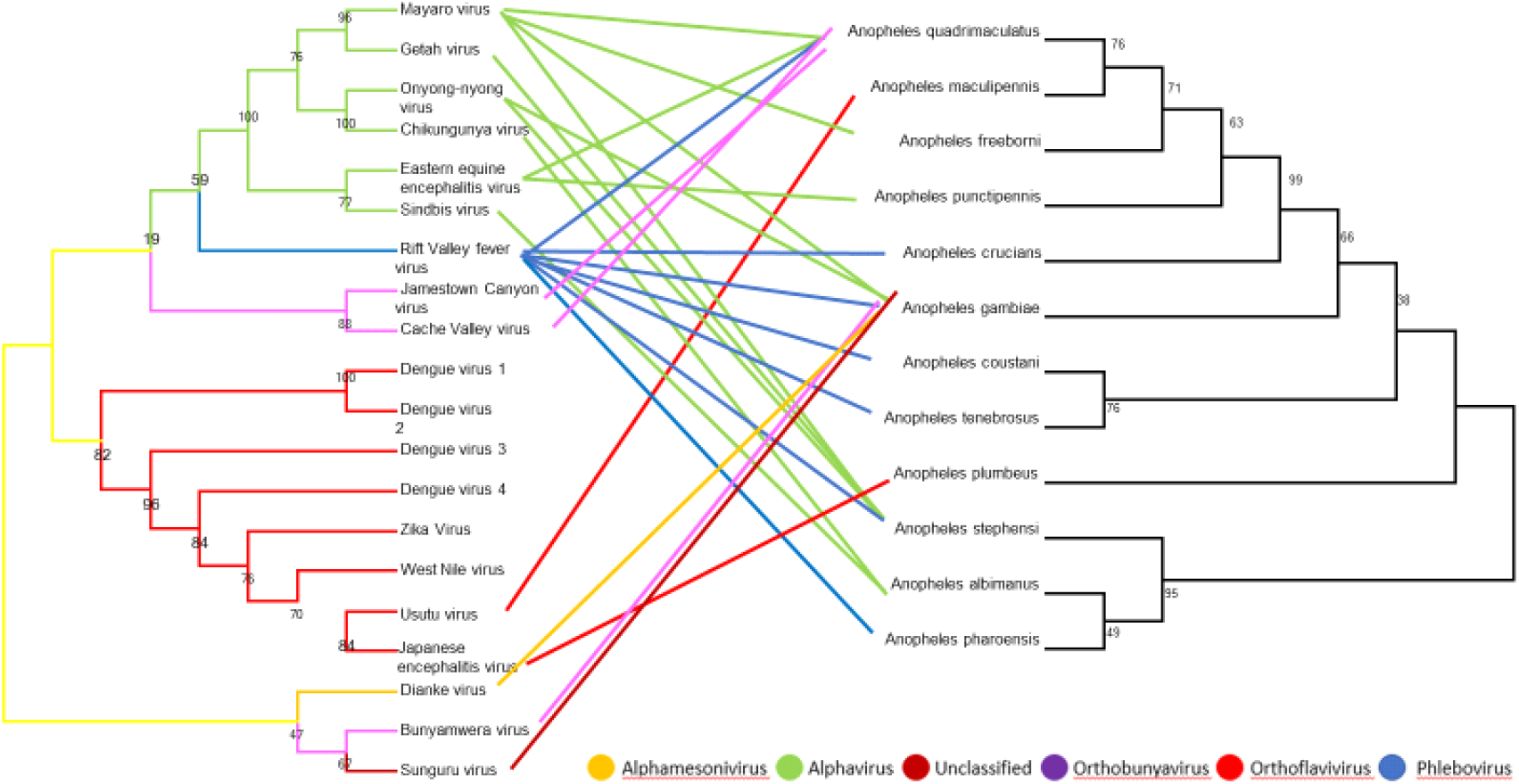
Tanglegram illustrating the phylogenetic relationships between *Anopheles* mosquito species (right tree, based on ITS2 genes) associated viruses (left tree, based on NS5 sequences). The phylogenies were inferred using the Neighbour-Joining method, incorporating sequences obtained from GenBank and original data. The **left panel** shows a phylogeny of 33 viruses, colour-coded by viral family: Alphavirus (green), Orthoflavivirus (red), Phlebovirus (blue), Orthobunyavirus (purple), Alphamesonivirus (yellow), and one unclassified virus (maroon). The **right panel** shows a phylogeny of 15 *Anopheles* mosquito species in which these viruses were experimentally tested. Coloured lines with bootstrap highlighted link each virus to the *Anopheles* species evaluated in laboratory competence studies.

**Figure 3.**
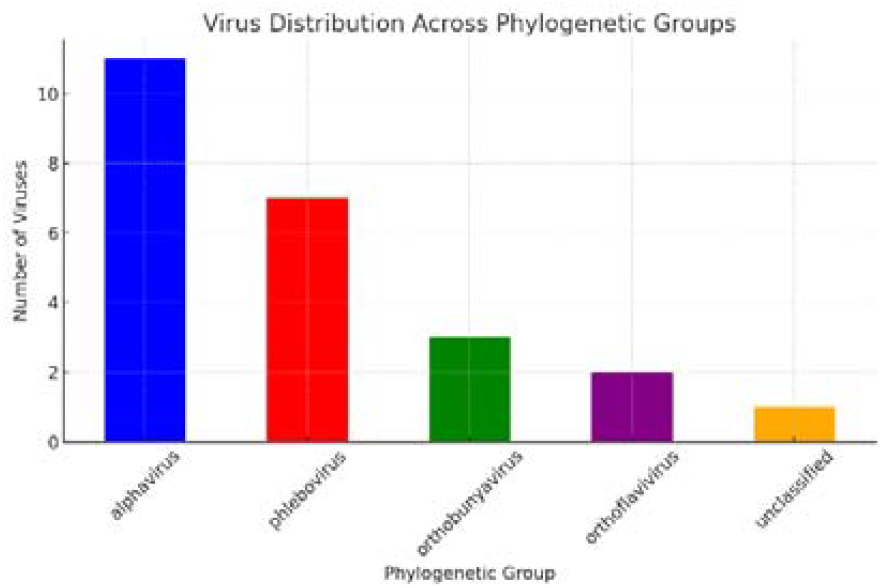
Distribution of viruses across phylogenetic groups, showing the number of viruses identified within each category. Alphaviruses are the most prevalent, followed by phleboviruses, while orthobunyaviruses, orthoflaviviruses, and unclassified viruses are less represented. This distribution suggests a higher occurrence or detectability of alphaviruses in the analyzed dataset.

**Figure 4.**
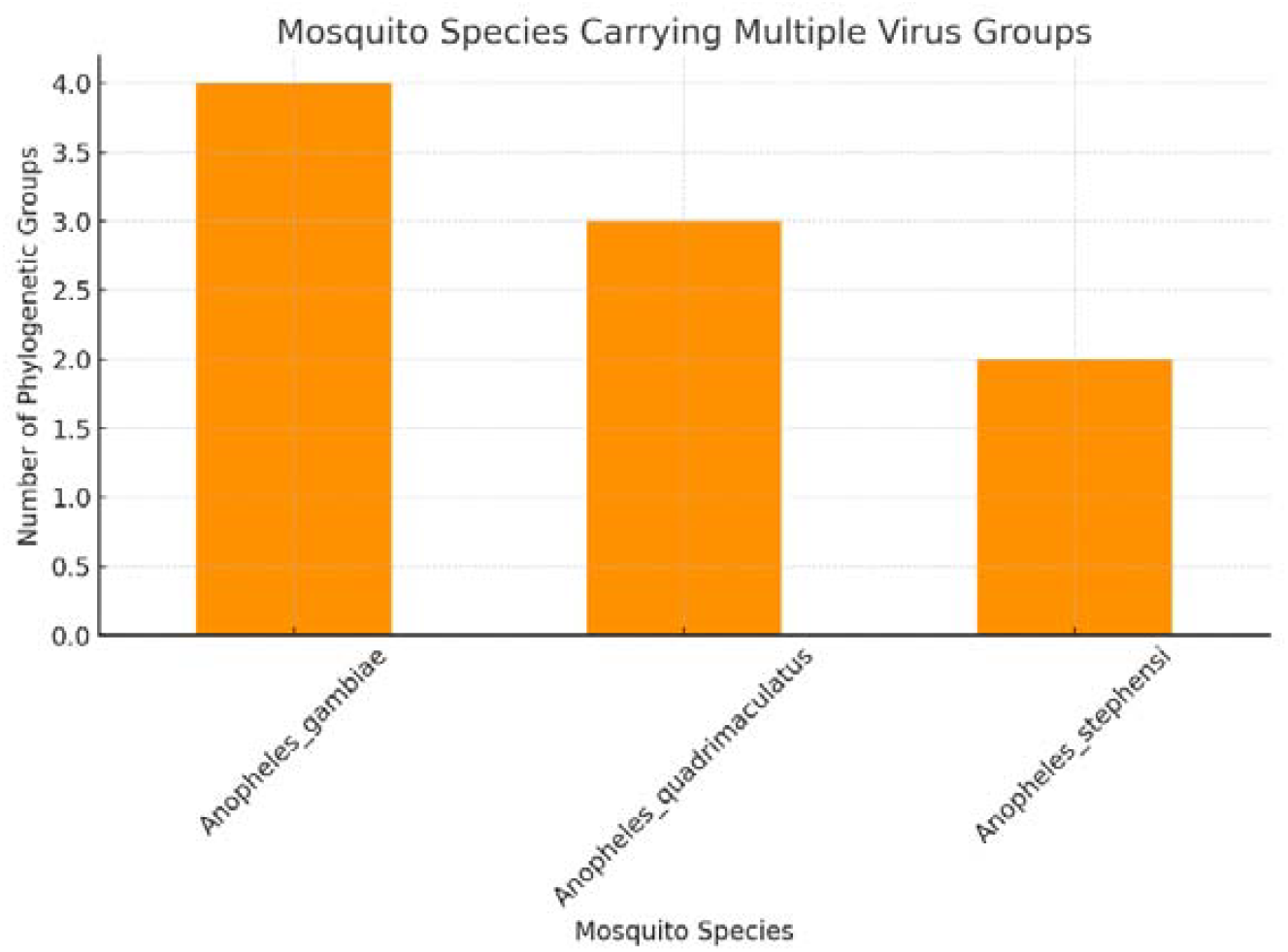
Number of phylogenetic virus groups detected in different *Anopheles* mosquito species. *Anopheles gambiae* harbors the highest number of virus groups (4), followed by *Anopheles quadrimaculatus* (3) and *Anopheles stephensi* (2). This suggests variation in the vector competence or exposure of different *Anopheles* species to diverse viral lineages.

These were tested across a wide array of Anopheles species, notably *An. gambiae, An. stephensi, An. quadrimaculatus*, and *An. albimanus*. The figure illustrates these frequent connections with multiple green edges linking alphaviruses to mosquitoes. However, the Orthoflaviviruses showed limited interactions with fewer mosquito species tested and mostly negative results, as seen in Table 1. These viruses were primarily tested *in An. stephensi, An. gambiae, and An. albimanus*. Phleboviruses, including Rift Valley fever virus, were tested in several African and U.S. Anopheles species, while Orthobunyaviruses and Alphamesoniviruses had more restricted representation.

Figure 2, most Importantly, also highlights taxonomic clustering of interactions certain clades of Anopheles, such as An. quadrimaculatus and its close relatives, were more commonly tested. The visual clustering of lines also suggests potential phylogenetic trends in host competence that warrant further investigation. Overall, Figure 2 underscores both the taxonomic bias in experimental data and the diverse potential of Anopheles mosquitoes to act as vectors for a wide range of arboviruses.

Moderately high infection rates (>40%) were noted for 11 viruses, SINV, RVFV, JEV, ONNV, NGAV, MAYV, GETV, EIV, EEEV, CVV, and BUNYV. However, from the 11, eight were able to transmit. RVFV and GETV showed a transmission rate of over 50%.

Where competence was found, the possible role in the transmission of viruses was further investigated by reviewing diapause studies (Table 2). *An. quadrimaculatus, An. Freeborni, An. macullipennis, An. puctipenis, An. crucians* and *An. plumbeus*, have all shown vector competence in both field and laboratory settings, and these species also diapause as adults.

**Table 2.**
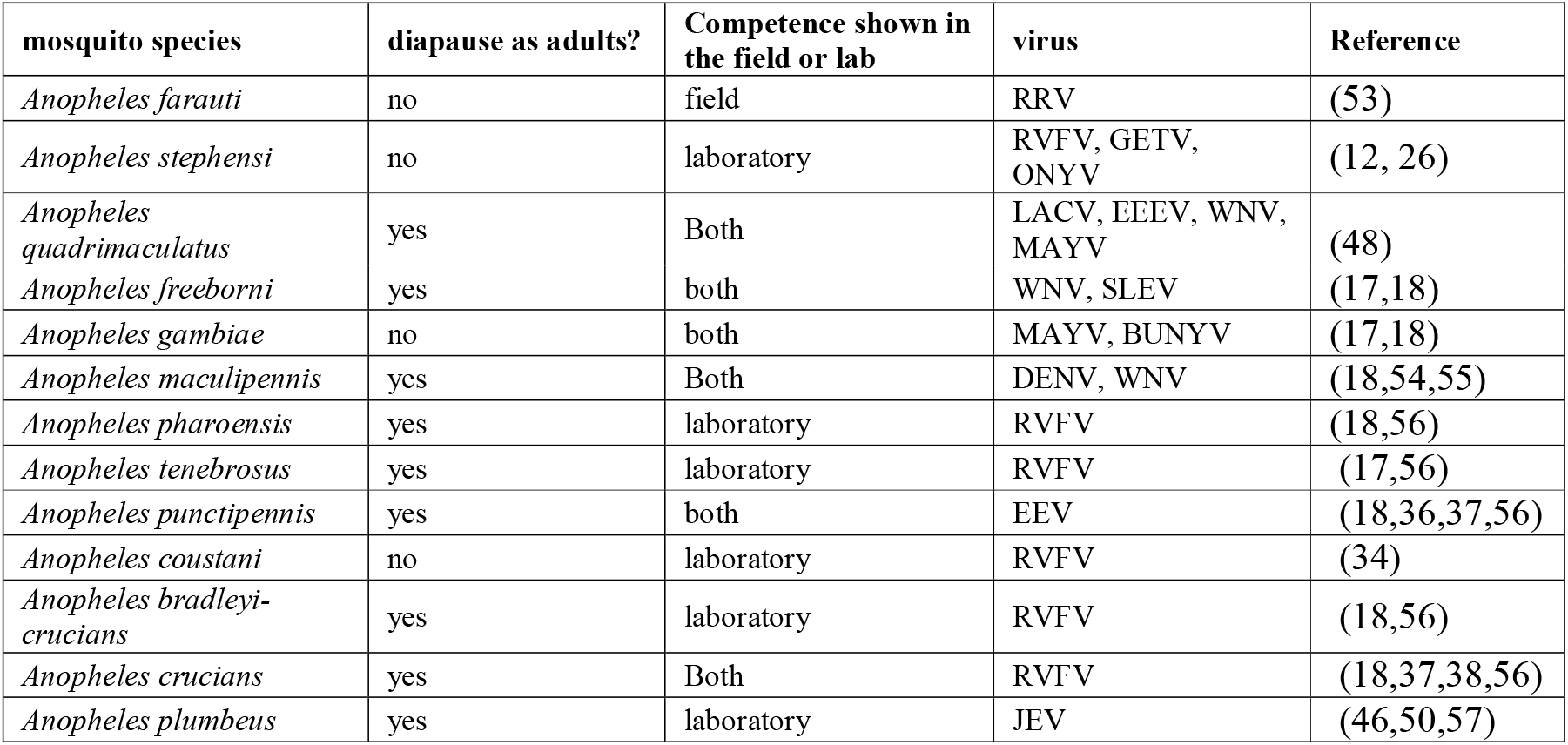
Mosquito diapause capability, vector competence demonstrated in laboratory or field settings, and the viruses for which they are competent, and references.

| mosquito species | diapause as adults? | Competence shown in the field or lab | virus | Reference |
| --- | --- | --- | --- | --- |
| *Anopheles farauti* | no | field | RRV | (53) |
| *Anopheles stephensi* | no | laboratory | RVFV, GETV, ONYV | (12, 26) |
| *Anopheles quadrimaculatus* | yes | Both | LACV, EEEV, WNV, MAYV | (48) |
| *Anopheles freeborni* | yes | both | WNV, SLEV | (17,18) |
| *Anopheles gambiae* | no | both | MAYV, BUNYV | (17,18) |
| *Anopheles maculipennis* | yes | Both | DENV, WNV | (18,54,55) |
| *Anopheles pharoensis* | yes | laboratory | RVFV | (18,56) |
| *Anopheles tenebrosus* | yes | laboratory | RVFV | (17,56) |
| *Anopheles punctipennis* | yes | both | EEV | (18,36,37,56) |
| *Anopheles coustani* | no | laboratory | RVFV | (34) |
| *Anopheles bradleyi-crucians* | yes | laboratory | RVFV | (18,56) |
| *Anopheles crucians* | yes | Both | RVFV | (18,37,38,56) |
| *Anopheles plumbeus* | yes | laboratory | JEV | (46,50,57) |

Comparative Phylogenetic Analysis of *Anopheles* Mosquitoes and Associated Viruses

## Discussion

While *Anopheles* mosquitoes are primarily known for malaria transmission, their potential role in arbovirus transmission, and as overwintering vectors, should not be overlooked. Comprehensive studies are required to understand the extent of this potential and its implications for public health. The results show that *Anopheles* mosquitoes are competent laboratory vectors to several viruses. We found that moderately high infection rates (> 40%) were noted for 11 viruses, SINV, RVFV, JEV, ONNV, NGAV, MAYV, GETV, EIV, EEEV, CVV, and BUNYV (table 1). In the results MAYV remains the most prevalent virus amongst most *Anopheles* species that have been tested, all tested *Anopheles* species were able to disseminate and transmit MAYV. RVFV was another common virus that showed high infection rates in *Anopheles* however, most studies did not continue to test dissemination or transmission. These findings highlight that *Anopheles* mosquitoes, though not primary vectors, may contribute to secondary transmission cycles in regions where environmental or ecological conditions limit the activity of primary vectors.

Previous field studies support this possibility. At least 51 different viruses have been detected in *Anopheles* spp, in the field (17). These studies have shown the potential to transmit diseases to both humans and other vertebrates. ONNV, SINV, RVFV, WNV. JEV and CVV are amongst some also seen in this study (17). Collectively, this evidence underscores that *Anopheles* mosquitoes are not merely incidental hosts, but may play ecologically meaningful roles in arbovirus maintenance.

A key finding from this review concerns the link between vector competence and diapause behaviour. Diapause allows mosquitoes to survive unfavourable environmental conditions by halting reproductive and metabolic processes (58). Our synthesis shows that several *Anopheles* species capable of adult diapause — including An. quadrimaculatus, *An. freeborni, An. maculipennis, An. punctipennis, An. crucians*, and *An. plumbeus* — have also demonstrated laboratory or field competence for one or more arboviruses. This dual capability suggests a plausible mechanism for viral overwintering.

For instance, *An. freeborni* exhibits adult diapause in low temperatures in California (12), while *An. maculipennis* has been observed overwintering in bunkers in the Netherlands (59), co-occurring with *Culex pipiens* mosquitoes infected with Usutu virus (USUV). In that study, USUV persisted in overwintering *Culex* species, suggesting that similar mechanisms could occur in diapausing *Anopheles* populations. Given that *An. maculipennis* has demonstrated high competence for WNV and DENV in laboratory studies (60), this raises the possibility that arboviruses could persist through winter within *Anopheles* populations in temperate zones.

This potential overwintering mechanism would allow transmission cycles to persist year-to-year, even when primary vector populations decline during cold seasons. Such persistence could have major epidemiological implications, particularly in temperate regions where viral activity typically resumes each summer.

Further experimental and modelling work is required to clarify the contribution of *Anopheles* species to arbovirus overwintering. For example, *An. stephensi*, which does not diapause, remains active year-round in certain regions and has demonstrated competence for JEV, GETV, RVFV, and ONNV (12, 26). Such species could act as continuous transmission reservoirs, whereas diapausing species might support virus persistence during inactive seasons.

Future modelling should incorporate the overwintering potential of *Anopheles* vectors into risk prediction frameworks. Doing so could extend the estimated range of arbovirus risk further polewards and increase the projected scale of annual outbreaks, as viruses may already be present in the environment when the transmission season begins, rather than requiring reintroduction each year.

## Conclusion

Our review demonstrates that *Anopheles* mosquitoes, traditionally viewed as vectors of Plasmodium parasites, are also experimentally competent for several arboviruses, including O’nyong-nyong virus (ONNV), Mayaro virus (MAYV), Rift Valley fever virus (RVFV), and others. Although competence levels are generally lower than in *Aedes* or *Culex* species, the evidence indicates that *Anopheles* mosquitoes can act as secondary or supporting vectors. Their potential contribution to virus maintenance, particularly through overwintering mechanisms, has likely been underestimated.

The association between vector competence and diapause suggests that *Anopheles* mosquitoes may play a previously unrecognised role in viral persistence during cold or dry seasons. Diapause enables adult mosquitoes to survive in temperate regions, potentially allowing virus survival between transmission seasons. Even if infection rates are low, the prolonged lifespan of diapausing mosquitoes could extend the window for virus maintenance and early-season re-emergence. Ultimately, recognising *Anopheles* mosquitoes as potential secondary arbovirus vectors adds a crucial dimension to vector surveillance and global disease control strategies.

## Notes

### Competing Interest Statement

The authors have declared no competing interest.

### Summary of Updates

Some minor changes in text and italicising some species

### Funding

- MRC Discovery Medicine North (DiMeN) Doctoral Training Partnership: MR/W006944/1

