## Supplementary material for "Evaluating the Role of *Anopheles* Mosquitoes in the Global Spread of Arboviruses: A Review of Laboratory-Confirmed Viral Competence": Table 1

Supplementary Materials

Supplementary material

Table 1. Data extracted from the *Anopheles’* data showing virus classification, mosquito species, Country of origin, vector competence and references. (supplementary material), low percentage values were defined as those ranging from 0% to 40%, whereas high percentage values were defined as those from 50% and above.

| **Virus Class** | **virus** | **mosquito species** | **mosquito origin (country)** | **Infection** | **Dissemination** | **Transmission** | **Reference** |
| --- | --- | --- | --- | --- | --- | --- | --- |
| Alphavirus | Getah virus | *Anopheles stephensi* | Japan | Yes | yes | yes | (21) |
| Orthoflavivirus | West Nile virus | *Anopheles farauti* | Australia | No | no | no | (22) |
| Alphavirus | Mayaro virus | *Anopheles albimanus* | United States | yes, high | yes | yes | (23) |
| Alphavirus | Sindbis virus | *Anopheles albimanus* | United States | yes, high | yes | yes | (23) |
| Alphavirus | Chikungunya virus | *Anopheles albimanus* | United States | yes, low | yes, low | yes, low | (23) |
| Orthoflavivirus | Dengue virus 2 | *Anopheles albimanus* | United States | No | not done | not done | (23) |
| Orthobunyavirus | Cache Valley virus | *Anopheles quadrimaculatus* | United States | yes high | yes | yes | (24) |
| Alphavirus | Mayaro virus | *Anopheles freeborni* | United States | yes, low | yes | yes | (25) |
| Alphavirus | Mayaro virus | *Anopheles gambiae* | United States | yes, high | yes | yes | (18) |
| Alphavirus | Mayaro virus | *Anopheles quadrimaculatus* | United States | yes, high | yes | yes | (25) |
| Alphavirus | Mayaro virus | *Anopheles stephensi* | United States | yes, high | yes | yes | (25) |
| Orthoflavivirus | Dengue virus 2 | *Anopheles maculipennis* | Italy | not done | no | no | (26) |
| Alphamesonivirus | Dianke virus | *Anopheles gambiae* | Senegal | yes, high | yes, low | no | (27) |
| Orthobunyavirus | Jamestown Canyon virus | *Anopheles quadrimaculatus* | United States | yes, high | yes, low | no | (28) |
| Orthoflavivirus | Zika virus | *Anopheles gambiae* | United States | no | no | no | (29) |
| Phlebovirus | Rift Valley fever virus | *Anopheles pharoensis* | Egypt | yes, high | yes, low | not done | (30) |
| Phlebovirus | Rift Valley fever virus | *Anopheles tenebrosus* | Egypt | yes, high | yes, low | not done | (30) |
| Alphavirus | Mayaro virus | *Anopheles quadrimaculatus* | United States | yes, high | yes, high | yes, low | (31) |
| Alphavirus | Eastern equine encephalitis virus | *Anopheles punctipennis* | United States | yes, high | not done | no | (32) |
| Alphavirus | Eastern equine encephalitis virus | *Anopheles quadrimaculatus* | United States | yes, high | not done | no | (32) |
| Phlebovirus | Rift Valley Fever virus | *Anopheles stephensi* | Netherlands | yes, low | yes, low | yes, low | (33) |
| Phlebovirus | Rift Valley fever virus | *Anopheles coustani* | Madagascar | yes, low | yes, low | yes, low | (34) |
| Phlebovirus | Rift Valley fever virus | *Anopheles gambiae* | Cameroon | not done | yes, low | not done | (35) |
| Phlebovirus | Rift Valley fever virus | *Anopheles bradleyi-crucians* | United States | yes, high | yes, low | not done | (36) |
| Phlebovirus | Rift Valley fever virus | *Anopheles crucians* | United States | yes, high | not done | not done | (37) |
| Phlebovirus | Rift Valley fever virus | *Anopheles quadrimaculatus* | United States | yes, high | no | no | (38) |
| Orthoflavivirus | Zika virus | *Anopheles quadrimaculatus* | United States | no | no | no | (39) |
| Sunrhavirus | Sunguru Virus | *Anopheles gambiae* | Uganda | yes, low | not done | not done | (40) |
| Orthobunyavirus | Bunyamwera virus | *Anopheles gambiae* | Kenya | yes, high | yes, high | not done | (41) |
| Orthobunyavirus | Cache Valley virus | *Anopheles quadrimaculatus* | United States | yes, high | yes, high | yes, low | (42) |
| Alphavirus | Onyong-nyong virus | *Anopheles stephensi* | Pakistan | yes, high | yes, high | yes, high | (43) |
| Orthoflavivirus | Usutu virus | *Anopheles plumbeus* | France | yes, low | yes, low | no | (44) |
| Alphavirus | Mayaro virus | *Anopheles gambiae* | NA | yes, high | yes, high | no | (45) |
| Orthoflavivirus | Japanese Encephalitis virus | *Anopheles plumbeus* | Belgium | yes, high | yes, low | yes, low | (46) |
| Alphavirus | Onyong-nyong virus | *Anopheles gambiae* | NA | yes, high | not done | not done | (47) |
| Alphavirus | Eastern equine encephalitis virus | *Anopheles quadrimaculatus* | United States | yes, high | yes, high | no | (48) |
| Alphavirus | Eilat virus | *Anopheles gambiae* | NA | yes, high | not done | no | (49) |
| Orthoflavivirus | Dengue virus 4 | *Anopheles stephensi* | Taiwan | no | not done | not done | (50) |
| Alphavirus | Onyong-nyong virus | *Anopheles gambiae* | NA | yes, high | not done | not done | (51) |
| Alphavirus | Eilat virus | *Anopheles gambiae* | NA | yes, high | yes, low | not done | (52) |

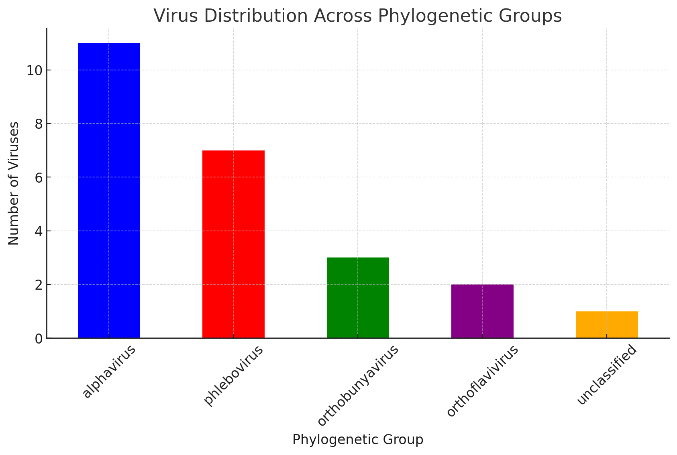

**Figure 3:** Distribution of viruses across phylogenetic groups, showing the number of viruses identified within each category. Alphaviruses are the most prevalent, followed by phleboviruses, while orthobunyaviruses, orthoflaviviruses, and unclassified viruses are less represented. This distribution suggests a higher occurrence or detectability of alphaviruses in the analyzed dataset.

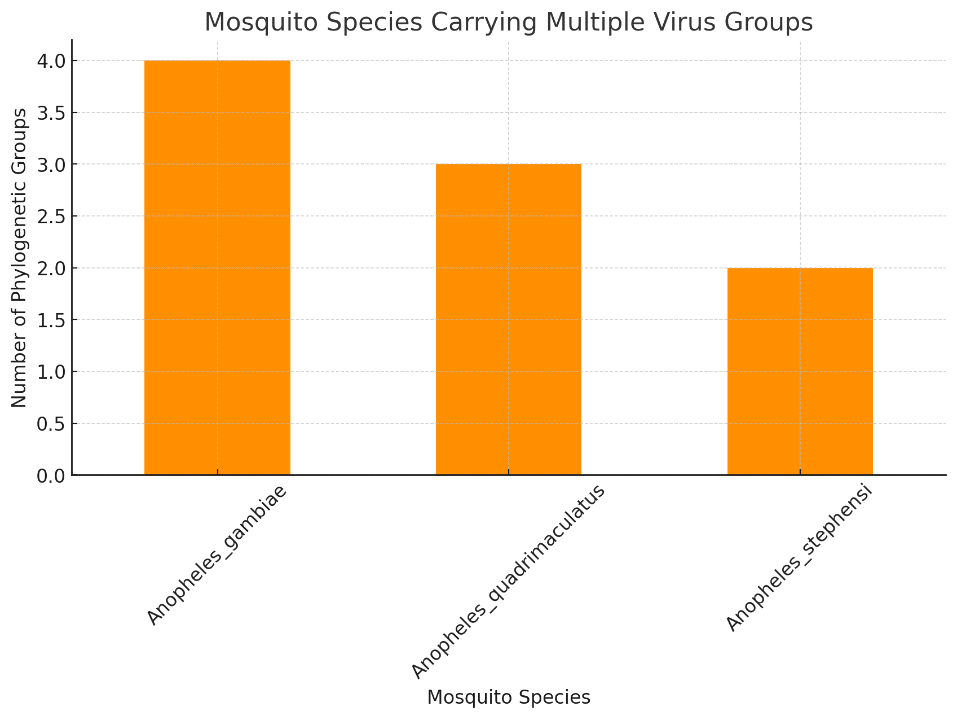

**Figure 4:** Number of phylogenetic virus groups detected in different *Anopheles* mosquito species. *Anopheles gambiae* harbors the highest number of virus groups (4), followed by *Anopheles quadrimaculatus* (3) and *Anopheles stephensi* (2). This suggests variation in the vector competence or exposure of different *Anopheles* species to diverse viral lineages.
